# Treemmer: a tool to reduce large phylogenetic datasets with minimal loss of diversity

**DOI:** 10.1101/249391

**Authors:** Fabrizio Menardo, Chloé Loiseau, Daniela Brites, Mireia Coscolla, Sebastian M. Gygli, Liliana K. Rutaihwa, Andrej Trauner, Christian Beisel, Sonia Borrell, Sebastien Gagneux

## Abstract

Large sequence datasets are difficult to visualize and handle. Additionally, they are often not an adequate representation of the natural diversity, but the result of uncoordinated and convenience sampling. Consequently, they can suffer from redundancy and sampling biases. Here we present Treemmer, a simple tool to evaluate the redundancy of phylogenetic trees and reduce their complexity by eliminating leaves that contribute the least to the tree diversity.

Treemmer can reduce the size of datasets with different phylogenetic structures and levels of redundancy while maintaining a sub-sample that is representative of the original diversity.

## Background

The number of genome sequences deposited into repositories such as NCBI and EBI is increasing rapidly. This wealth of data is at the same time a great opportunity and a challenge for biologists. Large datasets are difficult to visualize and use in downstream analyses. Additionally, being the product of different studies, downloaded datasets are often redundant and suffer from sampling biases. Several software packages such as CD-HIT [1] are available to reduce redundancy in a collection of amino-acid or nucleotide sequences [2]. Essentially, these methods cluster together sequences with a sequence identity higher than a certain threshold (specified by the user), and then select a representative sequence from each cluster for further analysis. While such methods are very efficient and can handle millions of sequences in a short time, they cannot be applied to whole genome data and do not consider phylogenetic relationships between sequences. To overcome these limitations, methods that reduce the size of datasets based on phylogenies instead of sequence similarity are needed. To date, two software packages have been developed for such purpose:

1) Tree pruner [3] is a tool to manually select and prune leaves/branches from a phylogenetic tree.
2) Treetrimmer [4] automatically reduces the number of leaves in a tree to few representatives for each user-defined operational taxonomical unit (OTU), like genus or species.

While both of these methods address the problem of size reduction in phylogenetic trees, they have some limitations: Tree pruner [3] can be very useful for manual curation, however it is not an automatic method, and it relies on subjective decisions by the users. Treetrimmer [4] is fully automatic, however it is based on user-defined OTU. For some datasets, information on the taxonomy might not be available and, more importantly, taxonomic categories are only a very rough proxy for genetic diversity.

Here we present Treemmer, a simple tool based on an iterative algorithm to reduce size and evaluate redundancy of phylogenetic datasets. In contrast to previous methods, Treemmer can automatically process any phylogenetic trees with branch lengths and does not require additional information. Treemmer prunes leaves from a phylogenetic tree while minimizing the loss of diversity. At each iteration, all pairs of neighboring leaves are evaluated, the pair with the shortest distance between leaves is selected, and one leaf is pruned off. The user can evaluate the redundancy of the dataset through the plot of the decay of the tree length and decide how many leaves to retain, or at what proportion of the original tree length to stop trimming. We applied Treemmer to two whole genome datasets (*Mycobacterium tuberculosis* and influenza A virus) and show that it can reduce their size and redundancy while maintaining a subset of samples that are representative of the overall diversity and topology of the original phylogenetic tree.

## Results

### The algorithm

We present here the algorithm implemented in Treemmer:

#### Step 1)

Given a phylogenetic tree, Treemmer iterates through all leaves, for each leaf it identifies the immediate neighboring leaves (separated by one node). If it does not find any immediate neighbor, it extends the search to leaves separated by two nodes. The result of this step is a list of pairs of neighboring leaves and their distances measured as the sum of the lengths of the branches separating the two leaves.

#### Step 2)

Treemmer selects the pair of leaves with the shortest distance among all the pairs of neighboring leaves, then it prunes a random leaf belonging to the pair. In case there are several equidistant pairs, Treemmer selects one at random. After pruning, all pairs of neighboring leaves containing the pruned leaf are eliminated from the list (Fig. 1).

**Figure 1.**
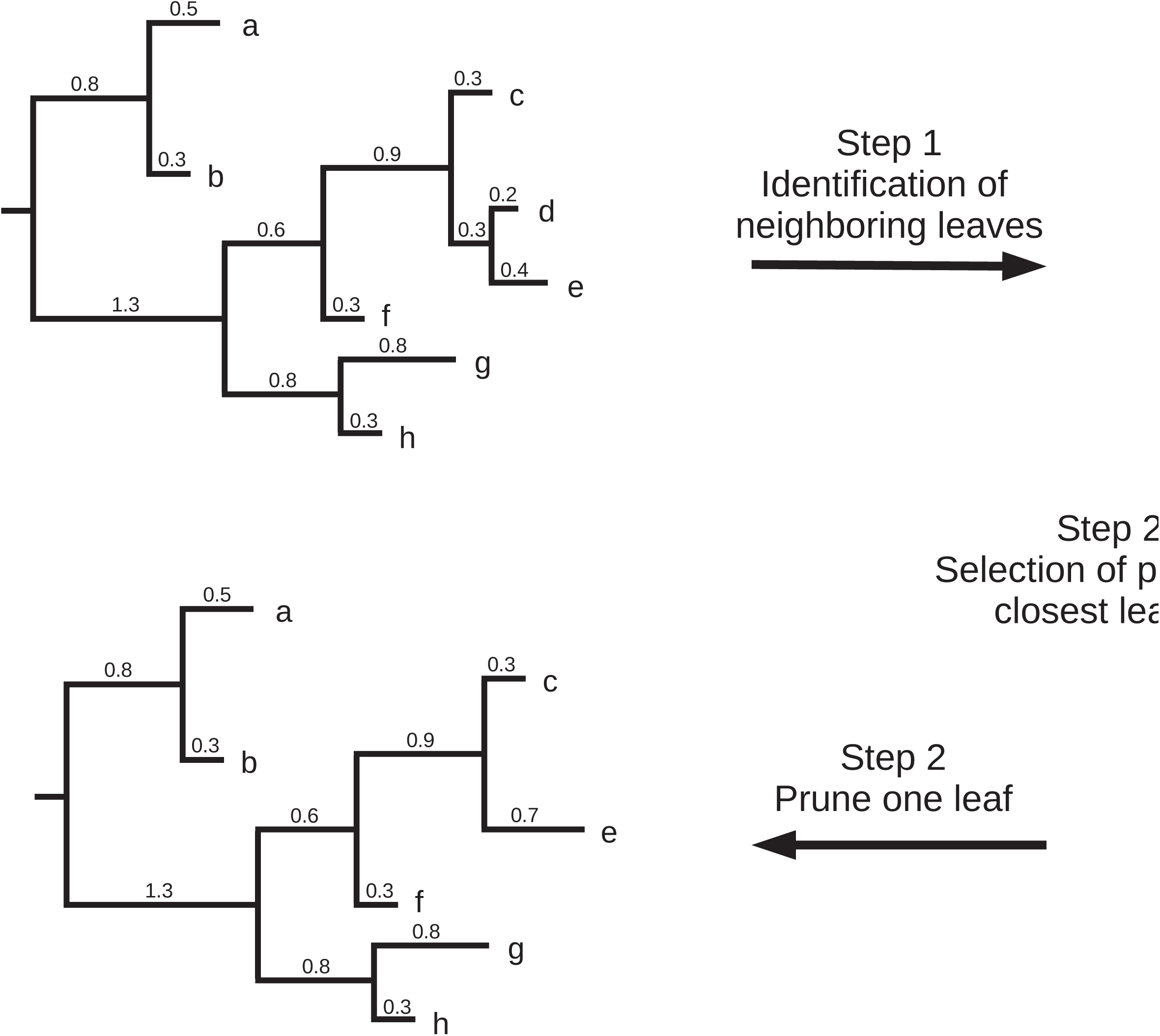
The core routine of Treemmer (with -*r* = 1): at each iteration the pair of closest leaves is identified and one of the two leaves is pruned from the tree, minimizing the loss of diversity.

Step 2 can be repeated any number of times before going back to step 1, this behavior can be controlled with the option -*r* (‐‐*resolution*). With the default value (-*r* = *1*), only one leaf is pruned (step 2) before the set of neighboring leaves is recalculated (step 1). With higher values of -*r* = x, step 2 is repeated x times (x leaves are pruned) before recalculating the set of neighboring leaves (step 1). Since step 1 is the most computationally intensive part of the algorithm, the option -*r* can speed up the running time of Treemmer considerably.

Steps 1 and 2 are repeated until one of the three possible stop criteria is reached:

#### Stop option 1)

The tree is pruned until there are three leaves left (default). At each iteration, the relative tree length (relative to the input tree) and the number of leaves in the tree are stored. Treemmer outputs the plot of the decay of the relative tree length (RTL, compared to the original tree). The RTL decay is a function of the redundancy of the dataset and of the phylogenetic structure of the tree. The RTL decay plot is a useful tool to evaluate the redundancy of the datasets and how much of the diversity is lost at each iteration.

#### Stop option 2)

With the stop option -*X* (‐‐*stop_at_X_leaves*) the user sets the number of leaves that should be retained in the reduced dataset. Treemmer prunes the input tree until the specified number of leaves is reached and outputs the list of retained leaves and the pruned tree. The output tree is useful for a quick evaluation of the reduced datasets but should not be used for further analysis, a new tree should be inferred with the reduced dataset.

#### Stop option 3)

With the stop option -*RTL* (‐‐*stop_at_RTL*) the user sets a threshold on the RTL of the reduced dataset. When the pruned tree RTL falls below the specified value, Treemmer stops pruning and outputs the list of retained leaves and the pruned tree. The output tree is useful for a quick evaluation of the reduced datasets but should not be used for further analysis, a new tree should be inferred with the reduced dataset.

### Polytomies

Treemmer can process trees with polytomies, however, very large unresolved polytomies (with thousands of leaves) increase considerably the number of pairs of neighboring leaves, slowing down the calculation. With the option -*p*/‐‐*solve-polytomies* all polytomies in the tree are resolved randomly with branch lengths set to zero

### Analysis of a *Mycobacterium tuberculosis* dataset

To demonstrate one possible application of Treemmer, we analyzed a *Mycobacterium tuberculosis* (MTB) tree built from the SNPs of 10,303 isolates. More information on the dataset and on the pipeline used to build the tree is available in the Methods section and in Supplementary Table 1.

This collection of genome sequences of MTB was not sub-sampled a priori, but represents all high-quality MTB genome sequences that we were able to retrieve from public repositories. This dataset is the result of several years of sampling by the scientific community, and while it covers most of the known diversity of MTB, it is highly redundant and suffers from sampling bias. One important source of redundancy originates from projects that analyzed individual MTB outbreaks with whole genome sequencing methods. In these projects, several identical or very similar strains were sequenced. Additionally, some lineages of MTB (i.e., L2 and L4) are overrepresented compared to others, mostly because they are predominant in countries where sampling was particularly extensive. We then tested whether Treemmer is able to reduce the redundancy and the sampling bias of this dataset.

We analyzed the decay of the relative tree length with four different values of -*r*: 1, 10, 100 and 1,000. We found that the RTL decay starts slowly and accelerates after about half of the leaves have been pruned. This confirms the high redundancy of this MTB dataset: pruning thousands of leaves affected the tree length only marginally, indicating that there were strains very similar to the pruned ones retained in the tree. Additionally, we found that the trajectories of the decay were overlapping and indistinguishable for -*r* = 1, -*r* = 10 and -*r* = 100. While for -*r* = 1,000, the decay was comparable to the other values of -*r* until the number of leaves reached about 6,000, and it was slightly faster with fewer leaves (Fig. 2). This finding suggests that the value of -*r* does not influence the results of Treemmer as long as it is two orders of magnitude smaller than the number of leaves. This is important for the analysis of large trees, because increasing the value of the -*r* option can considerably reduce the running time of Treemmer. While this result should be valid in general, we suggest, when possible, to start the analysis of new datasets running Treemmer with several different values of -*r* and comparing the results.

**Figure 2.**
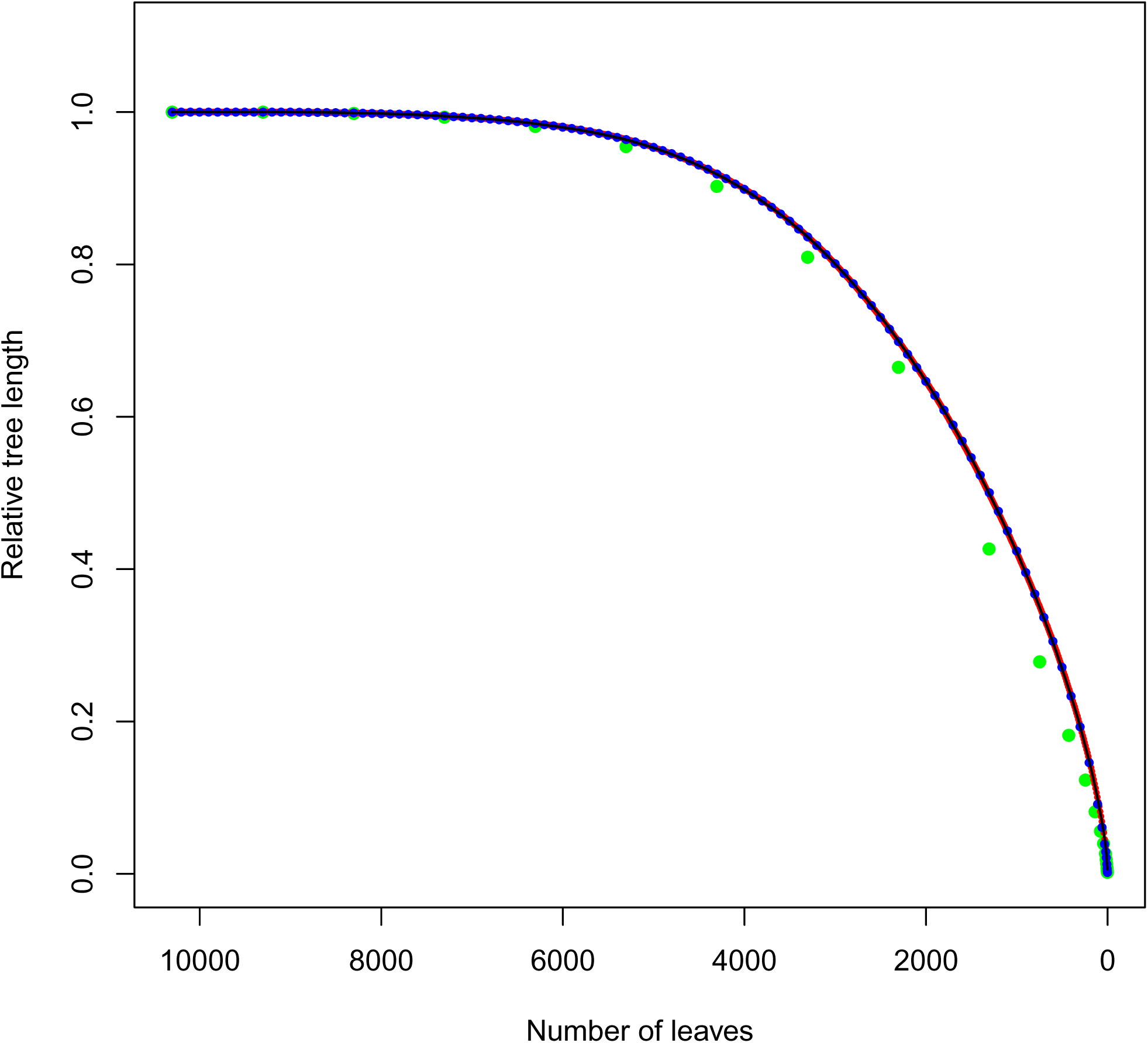
Plot of the relative tree length decay for the MTB dataset. Four different analysis were run with -*r* = 1 (black dots), -*r* = 10 (red dots), -*r* = 100 (blue dots) and -*r* = 1000 (green dots). The slow decay of the RTL is due to the high redundancy of the dataset. The RTL decays for -*r* = 1, 10 and 100 are overlapping and indistinguishable.

To obtain a reduced tree maintaining 95% of the original tree length, we ran Treemmer with the stop option -*RTL* 0.95. This resulted in a tree with 4,919 leaves, therefore Treemmer reduced the size of the dataset by more than 50 % while minimizing the loss of diversity (measured as tree length) to 5%. The resulting dataset has the same phylogenetic structure of the full-size original, is easier to handle, and can be used as a starting point for the downstream analysis (Fig. 3).

**Figure 3.**
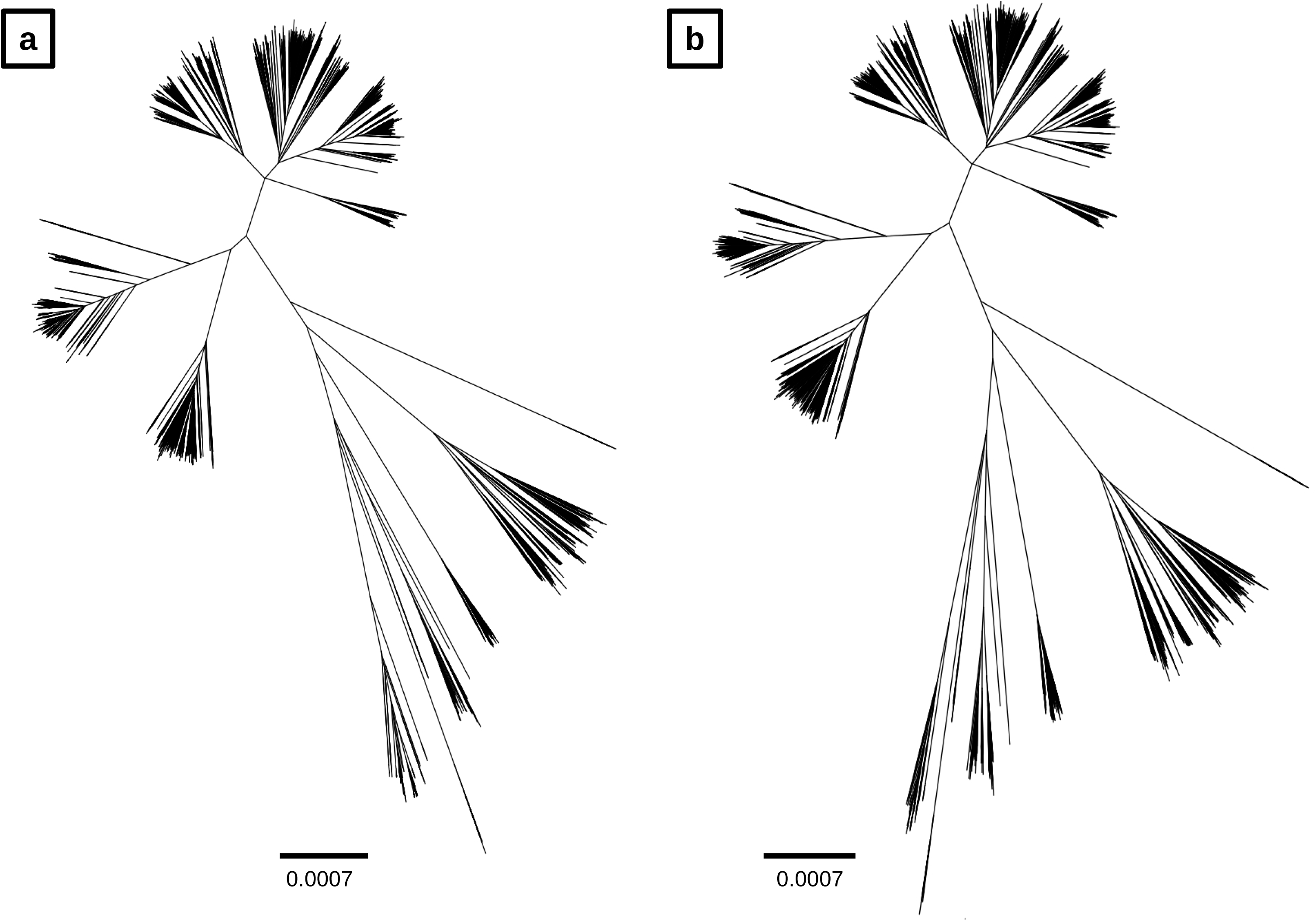
Comparison of original (a) and reduced (b) tree of the MTB dataset, with 10,303 and 4,919 leaves, respectively. The two trees have the same structure. The scale bar indicates expected substitution per position (only polymorphic nucleotide positions were included in the alignment).

### Analysis of the influenza dataset

We showed that Treemmer can reduce a redundant dataset while maximizing the retained diversity. Next, we tested Treemmer on a dataset with different characteristics and phylogenetic structure. To do this we used a time-calibrated influenza A / H3N2 tree with 2,080 viral sequences downloaded from nextstrain [5]. In contrast to the MTB dataset, this dataset is already a sub-sample of all the available influenza A genome sequences, filtered to reduce redundancy and to achieve an equitable temporal and geographic distribution [5]. Additionally, the influenza A tree has a different shape compared to the MTB tree: while the MTB tree comprises well-defined lineages that coalesce close to the root of the tree, the influenza tree is bushy and has a ladder-like structure (Fig. 4).

**Figure 4.**
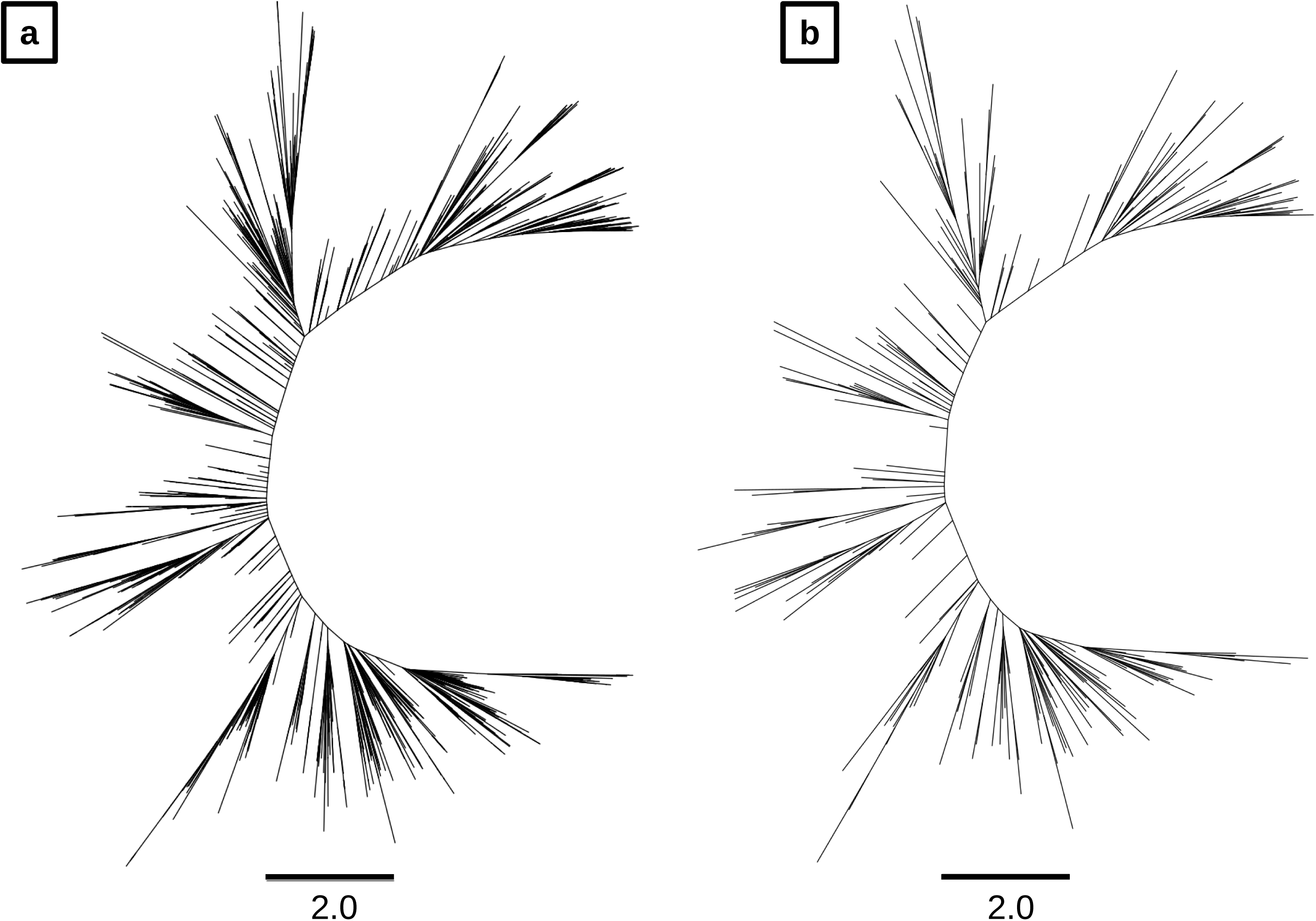
Comparison of original (a) and reduced (b) tree of the influenza A virus dataset, with 2,080 and 250 leaves, respectively. The scale bar indicates years.

We found that the trajectory of the RTL decay was steeper compared to the MTB dataset. The greater steepness of the RTL decay compared to the MTB tree is due to the reduced redundancy of the viral dataset: pruning few leaves reduces the tree length considerably. Additionally, we found that the decay of the relative tree length was not very different with three different values of -*r*: 1, 10 and 100 (Fig. 5). This confirms what we already observed during the analysis of the MTB dataset: the value of -*r* does not influence the results of Treemmer as long as it is smaller than two orders of magnitude compared to the number of leaves of the input tree.

**Figure 5.**
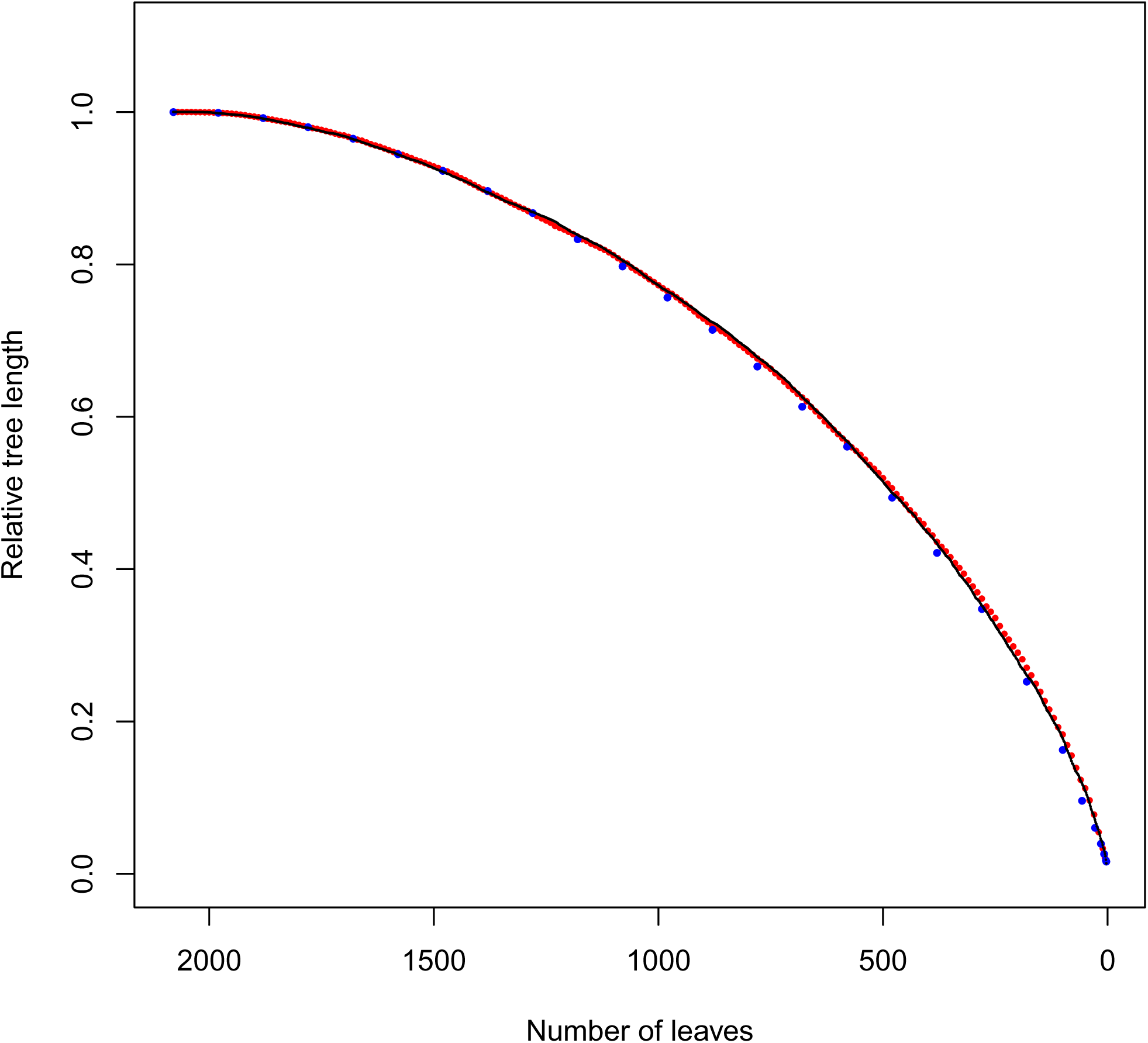
Plot of the relative tree length decay for the influenza A virus dataset. Three different analysis were run with -*r* = 1 (black dots), -*r* = 10 (red dots) and -*r* = 100 (blue dots). For this dataset the decay was faster than for the MTB dataset. This is due to the different structure of the phylogenetic trees and to the reduced redundancy of the viral dataset.

We then used Treemmer to subsample 250 leaves from the original influenza tree, corresponding to less than 40% of the original tree length. We found that the reduced tree has the same shape and phylogenetic structure of the original tree, demonstrating that Treemmer can be used to reduce the size of non-redundant datasets maintaining a representative set of samples (Fig. 4).

## Discussion

Many kinds of analyses, including bootstrap, tree calibration, fitting of a codon substitution model, or simply visualizing and browsing a phylogenetic tree, are impractical or impossible to perform with very large sequence datasets. To address these limitations, we developed Treemmer, a tool to reduce the size and redundancy of phylogenetic datasets while maintaining a representative diversity. In contrast to methods based on sequence similarity, Treemmer selects a representative subsample while considering the phylogenetic relationships between samples and their phylogenetic distance on the tree.

With the RTL decay plot it is possible to evaluate the loss of diversity when the size of the dataset is reduced. Treemmer can sample reduced datasets at any point of the RTL decay by selecting the balance between size and diversity that best fits the purpose of the user.

While the output of Treemmer can be used in many downstream analyses, it should not be considered as a random unbiased sample: the number of leaves belonging to different clades in the reduced dataset depends on the genetic diversity of the different clades and not on the abundance of different clades in nature; highly diverse clades will be represented by more leaves than less diverse ones, irrespectively of the frequency of such clades in natural populations. Additionally, some phenomena, e.g. recent fast speciation or population growth can result in large clades with short branches that would be pruned by Treemmer. Therefore, users interested in such phenomena should be careful when using Treemmer in their pipelines.

## Methods

### Implementation

Treemmer is written in python and uses the ETE library [6] to work with tree structures. Joblib [7] was used to parallelize the search of neighboring leaves (step1).

### *Mycobacterium tuberculosis* dataset

To assemble this dataset, we downloaded Illumina reads of 12,866 isolates of *M. tuberculosis* that we identified in the sequence read archive (SRA) repository. These represent the large majority of *M. tuberculosis* genomes currently available in the public domain.

Illumina adaptors were clipped and low quality reads were trimmed with Trimmomatic v 0.33 (SLIDINGWINDOW:5:20) [8]. Reads shorter than 20 bp were excluded for the downstream analysis. Overlapping paired-end reads were then merged with SeqPrep v 1.2 [9] (overlap size = 15). The resulting reads were mapped to the reconstructed ancestral sequence of the *M. tuberculosis* complex [10] with the mem algorithm of BWA v 0.7.13 [11]. Duplicated reads were marked by the MarkDuplicates module of Picard v 2.9.1 [12] The RealignerTargetCreator and IndelRealigner modules of GATK v 3.4.0 [13] were used to perform local realignment of reads around InDels. Pysam v 0.9.0 [14] was used to exclude reads with alignment score lower than (0.93*read_length)-(read_length*4*0.07)): this corresponds to more than 7 miss-matches per 100 bp. SNPs were called with Samtools v 1.2 mpileup [15] and VarScan v 2.4.1 [16] using the following thresholds: minimum mapping quality of 20, minimum base quality at a position of 20, minimum read depth at a position of 7X, minimum percentage of reads supporting the call 90%, maximum strand bias for a position 90%.

Strains with average coverage < 20 X were excluded. Additionally, we excluded genomes with more than 50 % of the SNPs excluded due to the strand bias filter. Furthermore, we excluded genomes with more than 50% of SNPs with a percentage of reads supporting the call included between 10% and 90%. Finally, we filtered out genomes with phylogenetic SNPs belonging to different lineages of MTB, as this is an indication that a mix of strains was sequenced.

After filtering, we obtained 338,553 positions with less than 10% of missing data were polymorphic in at least one strain. After eliminating strains with more than 10% of missing data at these positions, the final dataset comprised 10,303 strains with high-quality genomes (Supplementary Table 2).

The phylogenetic tree was inferred with FastTree [17] with options ‐nocat ‐nosupport and ‐fastest.

### Influenza A dataset

The influenza A virus tree was download from nextstrain [5] on the 1^st^ of December 2017. In the downloaded tree, few branches had small negative values, these were poorly supported branches that were collapsed before running Treemmer.

## Declarations

### Ethics approval and consent to participate

Not applicable.

### Consent for publication

Not applicable.

### Availability of data and materials

Treemmer is a free software under the GPL license, it is available together with the data used in this study here: https://git.scicore.unibas.ch/TBRU/Treemmer

### Authors’ contributions

FM wrote the software, FM, CL, DB, MC, SMG, LR, AT, SB and CB prepared the data and performed the analysis, FM and SG wrote the manuscript, all authors read and approved the final manuscript.

## Acknowledgments

The authors thank Richard Neher for his kind help with nextstrain. Calculations were performed at sciCORE (http://scicore.unibas.ch/) scientific computing core facility at the University of Basel.

## References

[1] Li, W., Jaroszewski, L., & Godzik, A. (2001). Clustering of highly homologous sequences to reduce the size of large protein databases. Bioinformatics, 17(3), 282–283.

[2] Sikic, K., & Carugo, O. (2010). Protein sequence redundancy reduction: comparison of various method. Bioinformation, 5(6), 234.

[3] Krishnamoorthy, M., Patel, P., Dimitrijevic, M., Dietrich, J., Green, M., & Macken, C. (2011). Tree pruner: An efficient tool for selecting data from a biased genetic database. BMC bioinformatics, 12(1), 51.

[4] Maruyama, S., Eveleigh, R. J., & Archibald, J. M. (2013). Treetrimmer: a method for phylogenetic dataset size reduction. BMC research notes, 6(1), 145.

[5] Neher, R. A., Bedford, T. (2015). nextflu: real-time tracking of seasonal influenza virus evolution in humans, Bioinformatics, 31(21) 3546–3548.

[6] Huerta-Cepas, J., Serra, F., & Bork, P. (2016). ETE 3: reconstruction, analysis, and visualization of phylogenomic data. Molecular biology and evolution, 33(6), 1635–1638.

[7] Joblib: https://pythonhosted.org/joblib.

[8] Bolger, A. M., Lohse, M., & Usadel, B. (2014). Trimmomatic: A flexible trimmer for Illumina. Sequence Data. Bioinformatics, 30(15), 2114–2120.

[9] SeqPrep: https://github.com/jstjohn/SeqPrep.

[10] Comas, I., Coscolla, M., Luo, T., Borrell, S., Holt, K. E., Kato-Maeda, M., … & S. Gagneux. (2013). Out-of-Africa migration and Neolithic coexpansion of Mycobacterium tuberculosis with modern humans. Nature genetics, 45(10), 1176–1182.

[11] Li H. and Durbin R. (2009) Fast and accurate short read alignment with Burrows-Wheeler Transform. Bioinformatics, 25:1754–60.

[12] Picard: https://github.com/broadinstitute/picard.

[13] McKenna A, Hanna M, Banks E, Sivachenko A, Cibulskis K, Kernytsky A, Garimella K, Altshuler D, Gabriel S, Daly M, DePristo MA, (2010). The Genome Analysis Toolkit: a MapReduce framework for analyzing next-generation DNA sequencing data. Genome Research, 20:1297–303.

[14] Pysam: https://github.com/pysam-developers/pysam.

[15] Li H. (2011). A statistical framework for SNP calling, mutation discovery, association mapping and population genetical parameter estimation from sequencing data. Bioinformatics, 27(21):2987–93.

[16] Koboldt, D., Zhang, Q., Larson, D., Shen, D., McLellan, M., Lin, L., Miller, C., Mardis, E., Ding, L., & Wilson, R. (2012). VarScan 2: Somatic mutation and copy number alteration discovery in cancer by exome sequencing Genome Research, 22(3), 568–576.

[17] Price, M.N., Dehal, P.S., and Arkin, A.P. (2010) FastTree 2 - Approximately Maximum-Likelihood Trees for Large Alignments. PLoS ONE, 5(3):e9490.

